# Effects of Variants of Concern Mutations on the Force-Stability of the SARS-CoV-2:ACE2 Interface and Virus Transmissibility

**DOI:** 10.1101/2023.01.06.522349

**Authors:** Magnus S. Bauer, Sophia Gruber, Adina Hausch, Marcelo C.R. Melo, Priscila S.F.C. Gomes, Thomas Nicolaus, Lukas F. Milles, Hermann E. Gaub, Rafael C. Bernardi, Jan Lipfert

## Abstract

Viruses mutate under a variety of selection pressures, allowing them to continuously adapt to their hosts. Mutations in SARS-CoV-2 have shown effective evasion of population immunity and increased affinity to host factors, in particular to the cellular receptor ACE2. However, in the dynamic environment of the respiratory tract forces act on the binding partners, which raises the question whether not only affinity, but also force-stability of the SARS-CoV-2:ACE2 bond, might be a selection factor for mutations. Here, we use magnetic tweezers (MT) to study the effect of amino acid substitutions in variants of concern (VOCs) on RBD:ACE2 bond kinetics with and without external load. We find higher affinity for all VOCs compared to wt, in good agreement with previous affinity measurements in bulk. In contrast, Alpha is the only VOC that shows significantly higher force stability compared to wt. Investigating the RBD:ACE2 interactions with molecular dynamics simulations, we are able to rationalize the mechanistic molecular origins of this increase in force-stability. Our study emphasizes the diversity of contributions to the assertiveness of variants and establishes force-stability as one of several factors for fitness. Understanding fitness-advantages opens the possibility for prediction of likely mutations allowing rapid adjustment of therapeutics, vaccination, and intervention measures.

## INTRODUCTION

Viruses constantly adapt to their hosts through genomic changes. While many mutations are silent or inviable, some result in increased fitness by increasing intrinsic transmissibility or evasion of population immunity. Better adaption and higher transmissibility enable new variants to supersede existing ones, as it was observed by the emergence and rapid spread of variants of concern (VOC) of SARS-CoV-2 (Supplementary Figure S1). Adaptations associated with increased fitness of SARS-CoV-2 often go along with higher affinity to host factors (*1, 2*). However, attachment of SARS-CoV-2 to the human host factor ACE2 takes place in the dynamic environment of the respiratory tract, where external forces, caused for example by breathing, coughing or mucus clearing by ciliary beating (*3, 4*), constantly counteract attachment (Figure 1A). In general, for successful infection, not only affinity, but also the stability under force for the attachment to the host can be critical (*5-11*). How force-stability varies across VOCs and whether it correlates with viral transmissibility is not well understood. Here, we use a highly sensitive single-molecule assay to mimic the natural force-exposure and investigate SARS-CoV-2 attachment to ACE2 under load. Our approach uses a fusion protein construct that features the receptor binding domain (RBD) from the SARS-CoV-2 spike protein coupled to the ACE2 ectodomain via a flexible peptide linker with molecular handles suitable for attachment in magnetic tweezers (MT; Figure 1B-D) (*12*). Coupling in MT enables us to probe the mechanical strength and dynamics of the RBD:ACE2 interaction under precisely defined forces (*12*) (Figure 1). Using our single-molecule assay, we characterize the interaction of SARS-CoV-2 wild type and VOC constructs with ACE2. Our assay reveals significant differences in force-stability between the VOCs and we identify the amino acid substitutions responsible for the changes. In addition to force-stability, we find that by extrapolating bond lifetimes to zero force, we can reproduce equilibrium affinity constants measured in bulk. Using molecular dynamics (MD) simulations, we identify key residues that stabilize the RBD:ACE2 interface and calculate the correlation between their movements. Comparing correlation coefficients for the different VOCs from MD simulations, we are able to accurately reproduce the experimentally determined relations in force-stabilities. Thus, we are able to experimentally determine differences in the force-stability between different VOCs using MT and elucidate their molecular origin with the help of MD simulations. Correlating our results to observations from epidemiology, we suggest that increased force-stability of the ACE2:RBD interface can increase intrinsic transmissibility. We anticipate that taking into account force-stability will aid predictions of future variants of concern.

**Figure 1:**
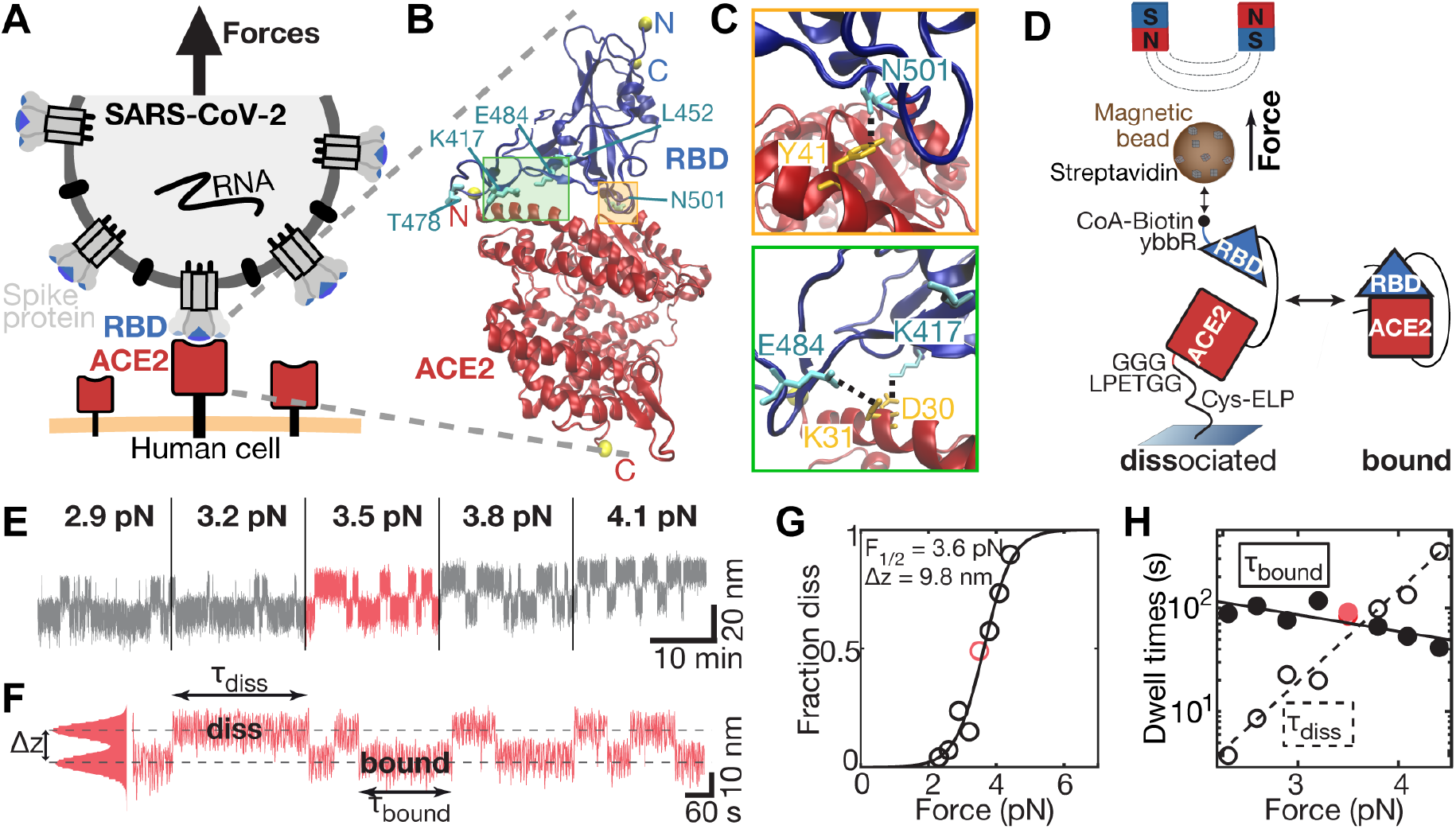
A single-molecule tethered ligand assay to study RBD binding to ACE2 for variants of concern of SARS-CoV-2. **A** A SARS-CoV-2 virion (grey) presents its spike protein trimers containing three RBDs (blue) ready for binding to human ACE2 (red). Attachment occurs in the dynamic environment of the respiratory tract, where the interaction must withstand external forces (black arrow). **B** Crystal structure of SARS-CoV-2 RBD (blue) bound to ACE2 (red) (PDB ID: 6m0j) (*1*). Termini of protein chains are marked with yellow spheres. Amino acid substitutions featured in current variants of concern (VOCs) are indicated in cyan. Crystal structure was rendered using VMD (*2*). **C** Zooms into the interface regions indicated in **B**. Top: RBD residue N501 featured in VOCs Alpha, Beta, and Gamma forms a hydrogen-bond with ACE2 residue Y41 (*1*). Bottom: RBD residues E484 and K417 featured in VOCs Beta and Gamma form salt bridges and hydrogen bonds with ACE2 residues K31 and D30 respectively (*1, 3*). Bridges and bonds are indicated as black dashed lines. **D** Schematic representation of the tethered ligand construct in MT. SARS-CoV-2 RBD (blue) is tethered via a flexible peptide linker to ACE2 (red). The construct is covalently attached to the glass surface via an ELP linker (see Materials and Methods), and to a streptavidin-covered magnetic bead via biotin at the C-terminus of the protein construct. **E** Representative extension time trace of the tethered ligand construct in MT shows binding and dissociation of the SARS-CoV-2 RBD:ACE2 interaction at plateaus of constant force. With increasing force, the interface is predominantly dissociated. **F** Segment of the trace in **E** at 3.5 pN, where it is equally distributed between the bound and dissociated (“diss”) state. The fraction dissociated at this force is thus 0.5. Dwell times in the dissociated state (τ_diss_) and in the bound state (τ_bound_) are indicated. **G** Fraction dissociated as a function of applied force (circles) and fit of a two-state model (solid line), with F_1/2_ and Δz (inset) as fitting parameters. **H** Mean dwell times in the bound and dissociated state for the molecule shown in **E**-**G** as a function of force (circles) and fit with an exponential model (solid lines).

## RESULTS AND DISCUSSION

### Comparing the force stability of the SARS-CoV-2 interaction with ACE2 for VOCs

We design fusion protein constructs comprising the ectodomain of the human ACE2 receptor connected by a flexible polypeptide linker to the RBD of SARS-CoV-2 for both the wild type (wt) strain and the Alpha, Beta, Gamma, and Delta VOCs (Figure 1C,D; Table 1). We attach the tethered ligand constructs to a glass surface and a magnetic bead, respectively, using a previously established protocol based on specific peptide linkers (*12-15*). Using MT, we probe the stability of the RBD:ACE2 interaction at varying levels of constant forces. At high forces larger than 25 pN, ACE2 unfolds with a characteristic unfolding signature (Supplementary Figure S2), enabling us to identify specific tethers. At forces below 10 pN, where both protein domains are folded, the linker ensures that receptor and ligand remain in proximity upon dissociation, allowing them to re-bind (*12, 16-18*). We can thus study repeated (re-)binding and dissociation of the same SARS-CoV-2 RBD and ACE2 interaction under different forces (Figure 1E). At low forces (< 2 pN), we find the bond to be predominantly formed, while increasing the force leads to elongated periods where the binding partners are dissociated. Quantitating the force dependence of the RBD:ACE2 interaction gives access to both the equilibrium force stability and the dynamics of the interactions.

**Table 1.** VOCs compared to SARS-CoV-2 wt. Number of molecules: 13 molecules (wt), 11 molecules (Alpha), 10 molecules (Beta), 14 molecules (Gamma), 20 molecules (Delta).

| WHO name | Pango lineage | Country of first observation | aa changes in RBD | $F_{1/2}$ (pN)<br>($F_{1/2}/F_{1/2}(\text{wt})$)<br>> 1: Higher force-stability than wt | $\tau_{0,\text{diss}}$ (s)<br>($t_{0,\text{diss}}/t_{0,\text{diss}}(\text{wt})$)<br>< 1: Faster binding than wt at $F = 0$ | $\tau_{0,\text{bound}}$ (s)<br>($t_{0,\text{bound}}/t_{0,\text{bound}}(\text{wt})$)<br>> 1: Slower dissociation than wt at $F = 0$ | $K_D$<br>($K_D/K_D(\text{wt})$)<br>< 1: Higher affinity than wt |
| --- | --- | --- | --- | --- | --- | --- | --- |
| wt | A | - | - | $3.8 \pm 0.4$<br>(1) | $0.06 \pm 0.17$<br>(1) | $121 \pm 284$<br>(1) | $532 \cdot 10^{-6}$<br>(1) |
| Alpha | B1.1.7 | UK | N501Y | $4.5 \pm 0.3$<br>(1.17) | $0.01 \pm 0.02$<br>(0.22) | $464 \pm 740$<br>(3.82) | $30 \cdot 10^{-6}$<br>(0.06) |
| Beta | B1.1.351 | South Africa | N501Y<br>E484K<br>K417N | $3.8 \pm 0.3$<br>(1.01) | $0.02 \pm 0.06$<br>(0.37) | $218 \pm 315$<br>(1.79) | $110 \cdot 10^{-6}$<br>(0.21) |
| Gamma | P1 | Brazil | N501Y<br>E484K<br>K417T | $4.0 \pm 0.6$<br>(1.06) | $0.02 \pm 0.04$<br>(0.27) | $552 \pm 1178$<br>(4.55) | $31 \cdot 10^{-6}$<br>(0.06) |
| Delta | B1.617.2 | India | L452R<br>T478K | $3.8 \pm 0.3$<br>(1.00) | $0.01 \pm 0.02$<br>(0.18) | $427 \pm 707$<br>(3.52) | $27 \cdot 10^{-6}$<br>(0.05) |

We characterize the force stability by *F*_1/2_, the midpoint force, at which it is equally likely to find the system in the bound or dissociated conformation (Figure 1F). We determine F_1/2_ by fitting a two-state model to the fraction dissociated (*f*_diss_) as a function of force (Figure 1G):

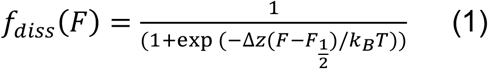

where *k*_B_ is Boltzmann’s constant and *T* the absolute temperature. *F*_1/2_ and Δz are fitting parameters that represent the midpoint force and the distance between the two states along the pulling direction, respectively. The fraction dissociated is calculated for each plateau of constant force by dividing the time spent in the dissociated conformation by the total plateau time. Comparing *F*_1/2_ for the wt and the different VOCs (Figure 2A,B and Supplementary Figure S3), we find that the force-stability for the Alpha VOC is highly significantly larger than wt (*p* = 5.2 · 10^−4^, two-tailed t-test). In contrast, we observe no statistically significant difference between SARS-CoV-2 wt and Beta (*p* > 0.80), wt and Gamma (*p* > 0.26), or wt and Delta (*p* > 0.99). In addition to the naturally emerged VOCs, we tested individual amino acid substitutions and found that E484K slightly lowers *F*_1/2_, albeit not statistically significantly, while K417N lowers the force stability highly significantly (*p* = 4.5 · 10^−5^).

**Figure 2:**
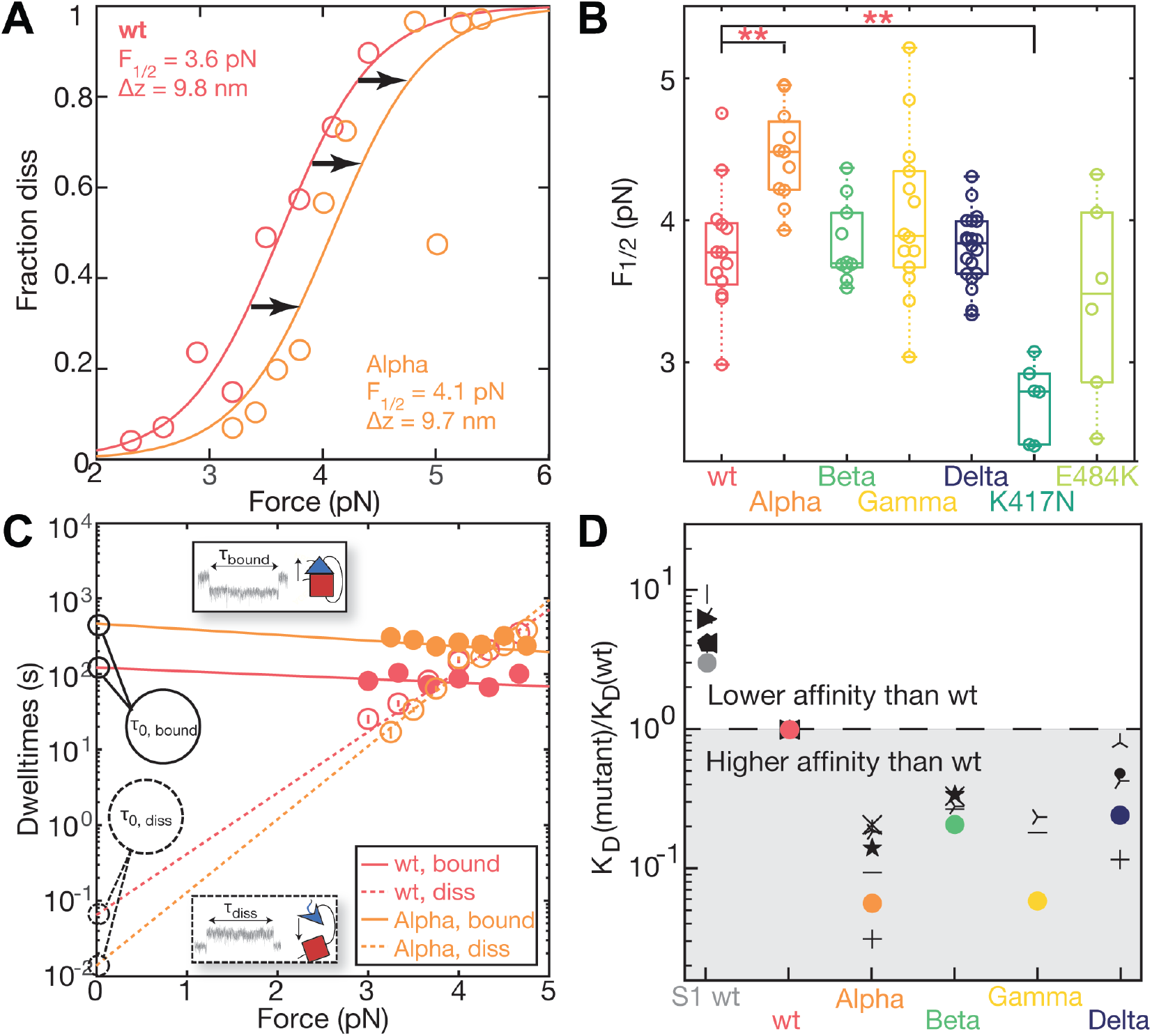
Effects of VOCs on interface force stability and affinity. **A** Representative force-dependent fraction dissociated for one wt (red), and one Alpha (orange) tethered ligand molecule. Points represent experimental data and solid lines two-state fits. **B** Midpoint forces determined for wt, VOCs, and K417N, and E484K single amino acid substitutions. F_1/2_ of Alpha (4.5 ± 0.33) pN and K417N (2.7 ± 0.27) pN deviate highly significantly from wt (3.8 ± 0.44) pN (p-values *p* = 0.00052 and *p* = 0.000045, respectively). Beta (3.8 ± 0.28) pN, Gamma (4.0 ± 0.57) pN, Delta (3.8 ± 0.25) pN, and E484K (3.4 ± 0.70) pN show no statistically significant difference to the wt. A bootstrap analysis revealed similar significance levels for the different variants (see Supplementary Figure S3). **C** Mean force-dependent dwell times (circles) and fit to the data in the bound (solid lines) and dissociated (dashed) state of wt and Alpha. Insets show example dwell times in MT traces. Additional comparisons for the other VOCs are in Supplementary Figure S4. **D** Mean dissociation constant normalized to the wt (K_D_/K_D_(wt); unnormalized K_D_s are in Supplementary Figure S5) determined from our measurements (colored dots) and compared to previous traditional bulk affinity measurements (SPR or BLI) (*3*–*9*) (Supplementary Table T2). Statistics in panels **B** - **C** reflect 13 molecules (wt), 11 molecules (Alpha), 10 molecules (Beta), 14 molecules (Gamma), 20 molecules (Delta), 6 molecules (E484K), and 6 molecules (K417N).

### Dynamics of the RBD:ACE2 interactions

To characterize the dynamics of the interactions, we determine the dwell times of the dissociated and bound state as a function of force. The dwell time distributions are well described by single exponentials (Supplementary Figure S4). The resulting mean dwell times, τ_diss_ and τ_bound_, (Figure 1H, circles) exhibit an exponential force dependence (Figure 1H, lines). The intersection of the fitted dwell times in the dissociated and bound state provides an alternative route to determine the midpoint force *F*_1/2_ and we find excellent agreement in both the absolute values and differences across VOCs with the *F*_1/2_ values determined using the two-state model (Equation 1; Supplementary Table T1). Further, extrapolation of the fits to zero force, assuming a slip bond behavior, yields lifetimes of the bond in the absence force (Figure 2C, Supplementary Table T1). The lifetimes of the dissociated state increase rapidly with increasing force for all constructs, which is expected as the peptide linker extends under force, which in turn impedes bond re-formation (*12, 16, 19, 20*). Conversely, the lifetimes of the bound state τ_bound_ decrease with force for all VOCs. However, the force dependence of τ_bound_ differs for the different VOCs (Supplementary Figure S5). For Alpha, we find a higher τ_bound_ than for wt over the whole measured force range (Figure 2C), again indicating a higher force-stability of Alpha in this force range, in line with observation of a higher *F*_1/2_ for Alpha. The other VOCs exhibit overall similar lifetimes in the bound state compared to wt under force, in line with their similar *F*_1/2_ values.

### Bond lifetimes at zero force predict affinities

The force-dependence of the bond lifetime (i.e. the slope in Figure 2C) varies considerably across the VOCs, leading to very different extrapolated lifetimes τ_0,bound_ and τ_0,diss_ in the absence of load. The lifetimes at zero force can be related to affinities: the ratio τ_0, diss_/τ_0, bound_ (or equivalently the ratio of the rates *k*_0,off_ / *k*_0,on_, which are the inverses of the lifetimes) define unimolecular equilibrium constants for the tethered ligand system *K*_D_^TL^ = τ_0,diss_/τ_0,bound_ (Supplementary Figure S6). The *K*_D_^TL^ are dimensionless, since both on- and off-rates for the tethered ligand systems are in units of 1/s. To compare *K*_D_^TL^ to bulk binding measurements, we need to take into account that in standard solution assays the bimolecular equilibrium constant *K*_D,sol_ = *k*_off,sol_ / *k*_on,sol_ has units of M, since while the off-rate *k*_off,sol_ is directly comparable to *k*_0,off_, the on-rate *k*_on,sol_ is depending on the ligand concentration, with unit of 1/M·s.

Therefore, we normalize both the unimolecular equilibrium constant *K*_D_^TL^ and the bimolecular dissociation constants *K*_D,sol_ obtained using standard binding assays (*1, 2, 21-25*) (*3*–*9*) to the wt and compare across the VOCs (Figure 2D). We find good agreement between the affinities determined in our single-molecule assay and the values obtained from standard affinity measurements (Figure 2D), suggesting that our constant force magnetic tweezers measurements are close enough to equilibrium to enable robust extrapolation to zero force. The good agreement between the force spectroscopy data extrapolated to zero force and the bulk binding measurements also implies that the RBD:ACE2 interaction is not a catch bond, but a “regular” slip bond. However, the data also reveal a clear difference between affinities at zero force and stability under mechanical load: While all VOCs exhibit increased affinity compared to wt, only Alpha shows higher force stability. A clear difference between force stability and thermodynamic affinity has been seen for a range of molecular bonds (*26, 27*) and our findings suggest that force stability must be taken into account as an independent factor when assessing VOCs.

### Molecular basis of different force stabilities across the VOCs

To provide a microscopic understanding of the observed differences in force stability, we performed all atom molecular dynamics (MD) simulations of the different RBD:ACE2 complexes. The structures of the VOCs were obtained by comparative modeling using Modeller (*28*), taking the SARS-CoV-2 wt structure as template (*12*). For each system, 5 replicas were simulated for a total of 1.0 μs using NAMD 3.0 (*29*). Trajectories were analyzed with VMD (*30*) and dynamical network analysis (*31*), which measures the correlation between the motions of neighboring residues to determine how cooperative their motion is. The higher the correlation between residues, the more relevant is their interaction for the stability of the protein complex (*32*). We determined contacts by tracking the distance between neighboring residues throughout the simulations, then selected the contacts with highest correlation for further analysis (see Supplementary Figure S7). Using these selection criteria, we identify the main contacts that are responsible for stabilizing the RBD:ACE2 complex (Figure 3A,B) and would thus be expected to have the greatest impact on the force-stability. All residues selected by the network analysis contact criterion are found within the receptor binding motif (RBM, residues N437 to Y508) region, at the RBD:ACE2 interface, defined as 4 Å cut-off based on the PDB ID 6m0j crystal structure (*33*) (see Supplementary Figure S8 for individual correlations). The total sum of correlations for RBD residues (Fig. 3C and Supporting Figure S9) highlights mostly hydrophobic residues and also residues that are not mutated in the VOCs, revealing that stabilizing the bond between RBD:ACE2 involves an interplay of different interactions. Mutations in the VOCs lead to a rebalancing of interactions with ACE2, losing correlation in some regions while gaining in others (Figure 3C and Supplementary Figure S7).

**Figure 3:**
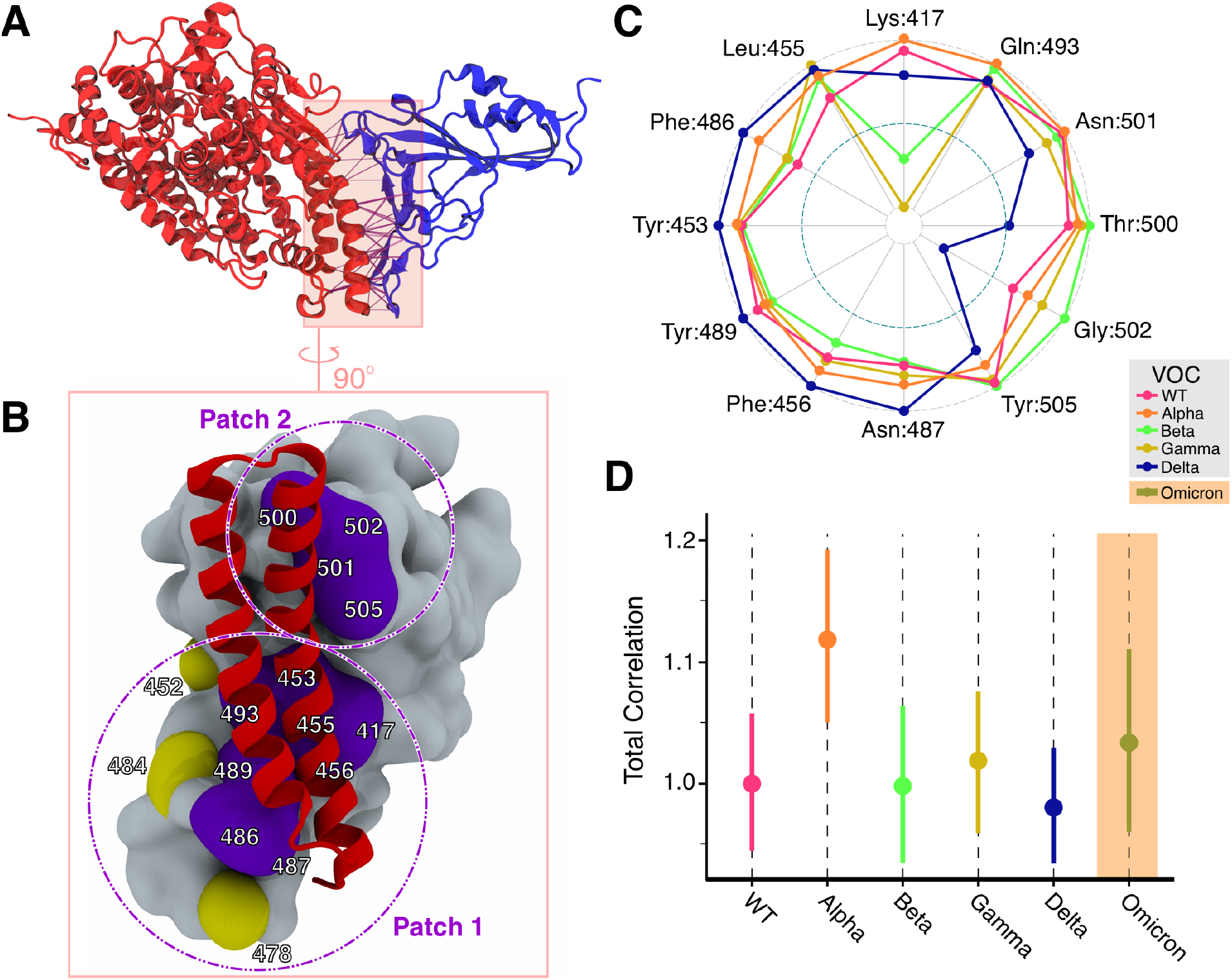
Effects of VOCs on the contact network at RBD:ACE2 interface. **A** Network interactions between Spike RBD (blue) complexed with ACE2 (red) represented as purple dashed cylinders, where thicker cylinders represents highly connected residues. **B** Detailed RBD:ACE2 surface interface highlighting the receptor binding domain (RBD) in grey. Residues colored in violet are highly correlated while residues colored in yellow are less correlated. **C** Total correlations for selected residues for the different VOCs. Values are normalized to the VOC with the strongest correlation. **D** Total correlation of motion summarized from the dynamic network interactions of all main correlations across the RBD:ACE2 interface, for each VOC. Total correlation measurements for all variants were normalized by the mean total correlation in the WT. Bars represent the 90% confidence intervals for the mean.

Examining the specific mutations involved in the VOCs, we can rationalize the observed differences in force stability. Alpha carries only one substitution in the RBD, namely N501Y, which is also present in Beta and Gamma. The N501Y substitution creates an additional hydrogen bond and leads to increased pi-stacking with residue Y41 of ACE2 (*34*), which increases the pairwise correlation (Supplementary Figure S8 and S9) and in particular enhances the correlations in the vicinity of position N501, in particular at residue T500 and G502 (Figure 3C).

The Beta and Gamma VOCs feature, in addition to N501Y, amino acid substitutions at position K417N/T and E484K. In the wt, the residues K417 and E484 form salt bridges with residues D30 and K31 of ACE2, respectively (*1, 3*). Due to a charge-reversal in the case of E484K and a charge removal in the case of K417N (Beta) and K417T (Gamma), these salt bridges are disrupted. The disruption of the salt bridge at position 417 leads to dramatic loss in correlation (Figure 3C), fully in line with a dramatically decreased force stability for K417N (Figure 2B). The charge reversal by the mutation E484K leads to the loss of the salt bridge with K31, which is, however, compensated by an alternative salt bridge with E35 (*3*). As a result, the correlations around position E484 are virtually unaffected by the mutation (see e.g. residues F486, N487, Y489; Figure 3C), consistent with only a slight and statistically insignificant decrease in *F*_1/2_ for the E484K mutation (Figure 2B). In summary, for the Beta and Gamma VOCs, the increased force stability by N501Y is offset by the reduction of stability in particular by K417N/T, such that the total effect is a similar force stability as wt.

The mutations in the Delta VOC, L452R and T478K, are located further away from the RBD:ACE2 interface and there are no direct interactions involving these residues. Nonetheless, the Delta mutations do shift the correlation pattern, with correlations in the “Patch 2” region (Figure 3B) around position N501 decreasing and increasing in the region “Patch 1” around P479.

To summarize the contact network and the correlations of motion in a single metric and to directly compare to the experimental force stability measurements, we define the total correlation as the sum of all correlations between stable contacts identified through the network analysis. The wt, the experimentally examined VOCs, and also the more recently emerged Omicron VOC were simulated and total correlation scores were computed, using a bootstrapping approach to determine confidence intervals (Figure 3D and Supplementary Figure S10). We find excellent agreement between the total correlation metric and the experimentally determined values for the force stability assessed by *F*_1/2_: wt, Beta, Gamma, and Delta exhibit the same force stabilities, while Alpha has a statistically larger *F*_1/2_ and total correlation. Interestingly, the recent Omicron variant, which features a very large number of mutations (15 in the RBD alone; (*35*)), has a total correlation value between Alpha and the wt, and did not show statistically significant differences when compared to Alpha and wt.

## Discussion

We have used our tethered ligand assay and exploited the high sensitivity of MT and their ability to measure at constant forces to precisely determine both the force stability and affinity of different SARS-CoV-2 VOCs binding to ACE2. Unlike traditional bulk affinity measurements, our assay provides additional insights into bond-stability and kinetics under mechanical load, mimicking the natural binding circumstances in the dynamic environment of the respiratory tract. We find that while the Alpha, Beta, Gamma, and Delta VOCs all have increased affinities to ACE2 compared to wt, only Alpha has higher force stability. Viral fitness depends on multiple factors, including cleavage and cell entry, replication, and viral packing and release (*36-41*). Correspondingly, the VOCs carry mutations in positions beyond the RBD. However, attachment to the host cell is a critical first step in viral infection and it is instructive to relate trends observed epidemiologically with our findings about the RBD:ACE2 interaction. The Alpha, Beta, and Gamma VOCs variants all emerged in the second half of 2020 and began to circulate widely in late 2020 and early 2021 (Supplementary Figure S1). In Europe and Northern America, Alpha, but not Beta or Gamma, quickly replaced the wt and became the dominantly circulating variant (Supplementary Figure S1). At that time, there was no widespread population immunity in Northern America or Europe (Supplementary Figure S1), suggesting that Alpha, but not Beta or Gamma, has a significant advantage in a (largely) immune-naïve population. We speculate that the higher force stability of Alpha contributes to its epidemiologically observed higher transmissibility compared to wt, Beta, or Gamma. The Beta and Gamma variants became dominant in South Africa and Brazil (Supplementary Figure S1), in a setting where there was already significant population immunity through natural infections (*42, 43*). The charge change mutations at positions K417 and E484 present in Beta and Gamma reduce the binding of certain neutralizing antibodies (*44*), but lower the force stability. Consequently, the Beta and Gamma appear to confer an advantage to the virus in population with significant immunity, but not in an immune-naïve population. The Delta and Omicron VOCs became dominant globally in mid 2021 and early 2022, respectively, at a time where there was significant population immunity through both vaccinations and natural infections (Supplementary Figure S1). Both variants show immune escape (*25, 45-47*) and likely enhanced transmission through substitution at the S1/S2 cleavage site, yet no, or only slightly increased force stability (Figure 2B and Figure 3D) (*48-50*). These observations raise the possibility that a new variant might emerge that combines the fitness advantages present in Omicron with a higher force stability, which could make it even more transmissible. Our correlation based analysis of the RBD:ACE2 interface by MD simulations and high-resolution force spectroscopy measurements of the interface in the tethered ligand assay has the potential to help understand and ultimately predict the spread of SARS-CoV-2 variants.

## ACKNOWLEDGMENTS

We thank Meike Bos, Joost de Graf, and David Dulin for helpful discussions and Nina Beier, Benedikt Böck, and Ellis Durner for help with initial experiments. This study was supported by German Research Foundation Projects 386143268 and 111166240, a Human Frontier Science Program Cross Disciplinary Fellowship (LT000395/2020C) and European Molecular Biology Organization Non-Stipendiary long-term fellowship (ALTF 1047-2019) to L.F.M., and the Physics Department of LMU Munich. R.C.B. and P.S.F.C.G. are supported by start-up funds provided by Auburn University.

