## Supplementary Information for "Effects of Variants of Concern Mutations on the Force-Stability of the SARS-CoV-2:ACE2 Interface and Virus Transmissibility"

**Content:**

**Supplementary Materials and Methods**
**Full Protein Sequences**
**Supplementary Table T1 – T2**
**Supplementary Figures S1 – S10**
**Supplementary References**

### SUPPLEMENTARY MATERIALS AND METHODS

All chemicals used were supplied by Carl Roth (Karlsruhe, Germany) or Sigma-Aldrich (St. Louis, MO, USA) unless otherwise noted.

#### Cloning and protein construct design

Tethered-ligand fusion proteins were created, expressed, and purified as previously described (*1*). Constructs for ACE2-linker-RBD of SARS-CoV-1 were designed in SnapGene Version 4.2.11 (GSL Biotech LLC, San Diego, CA, USA) based on a combination of the ACE2 sequence from Komatsu et al.(*2*) available from GenBank under accession number AB046569 and the SARS-CoV-1 sequence from Marra et al.(*3*) available from GenBank under accession number AY274119. The crystal structure by Li et al. (*4*) available from the Protein Data Bank (PDB ID: 2ajf) was used as a reference. The linker sequence and tag placement were adapted from Milles *et al.* (*5*). The linker sequence is a combination of two sequences available at the iGEM parts databank (accession numbers BBa_K404300, BBa_K243029). The fusion protein with the sequence of the RBD of SARS-CoV-2 was designed from the sequence published by Wu et al. (*6*) available from GenBank under accession number MN908947, with a 6x histidine (His) tag added for purification. In addition, tags for specific pulling in magnetic tweezers and the atomic force microscope were introduced: a triple glycine for sortase-mediated attachment on the N-terminus and a ybbR-tag, AviTag, and Fgɣ tag on the C-terminus. In summary, the basic construct is built up as follows: MGGG-ACE2-linker-RBD-6xHIS-ybbR-AviTag-Fgɣ. All protein sequences are provided in the full protein sequences paragraph.

The constructs were cloned using Gibson assembly from linear DNA fragments (GeneArt, ThermoFisher Scientific, Regensburg, Germany) containing the sequence of choice codon-optimized for expression in *E. coli* into a Thermo Scientific pT7CFE1-NHis-GST-CHA Vector (Product No. 88871). The mutations found in variants of concern causing amino acid substitutions in the RBD were introduced by blunt-end cloning and ligation. Replication of DNA plasmids was obtained by transforming in DH5-Alpha Cells and running overnight cultures with 7 ml lysogeny broth with 50 µg/ml carbenicillin. Plasmids were harvested using a QIAprep® Spin Miniprep Kit (QIAGEN, Germantown, MD, USA, # 27106).

#### In vitro protein expression

Expression was conducted according to the manual of 1-Step Human High-Yield Mini in vitro translation (IVT) kit (# 88891X) distributed by ThermoFisher Scientific (Pierce Biotechnology, Rockford, IL, USA). All components, except 5X dialysis buffer, were thawed on ice until completely thawed. 5X dialysis buffer was thawed for 15 minutes and 280 µl were diluted into 1120 µl nuclease-free water to obtain a 1X dialysis buffer. The dialysis device provided was placed into the dialysis buffer and kept at room temperature until it was filled with the expression mix.

For preparing the IVT expression mix, 50 µl of the HeLa lysate was mixed with 10 µl of accessory proteins. After each pipetting step, the solution was gently mixed by stirring with the pipette. Then the HeLa lysate and accessory proteins mix was incubated for 10 minutes. Afterwards, 20 µl of the reaction mix was added. Then 8 µl of the specifically cloned DNA (0.5 µg/µl) was added. The reaction mix was then topped off with 12 µl of nuclease-free water to obtain a total of 100 µl. This mix was briefly centrifuged at 10,000 g for 2 minutes. A small white pellet appeared. The supernatant was filled into the dialysis device placed in the 1X dialysis buffer. The entire reaction was then incubated for 16 h at 30°C under constant shaking at 700 rpm. For incubation and shaking a ThermoMixer comfort 5355 (Eppendorf AG, Hamburg, Germany, # 5355) with a 2 ml insert was used. After 16 h the expression mix was removed and stored in a protein low binding reaction tube on ice until further use.

**Protein purification**

Purification was conducted using HIS Mag Sepharose® Excel beads (Cytiva Europe GmbH, Freiburg, Germany, # 17371222) together with a MagRack™ 6 (Cytiva Europe GmbH, Freiburg, Germany, # 28948964) following the vendor's protocol. Bead slurry was mixed thoroughly by vortexing. 200 µl of homogenous beads were dispersed in a 1.5 ml protein low binding reaction tube. Afterwards the reaction tube was placed in the magnetic rack and the stock buffer was removed. Next, the beads were washed with 500 µl of HIS wash buffer (25 mM TRIS-HCl, 300 mM NaCl, 20 mM imidazole, 10% vol. glycerol, 0.25 % vol. Tween 20, pH 7.8). Expressed protein from IVTT was filled to 1000 µl with TRIS buffered saline (25 mM TRIS, 72 mM NaCl, 1 mM $CaCl_2$, pH 7.2) and mixed with freshly washed beads. The mix was incubated in a shaker for 1 h at room temperature. Subsequently, the reaction tube was placed in the magnetic rack and the liquid was removed. The beads were washed three times with wash buffer, keeping the total incubation time to less than 1 min. Remaining wash buffer was removed and 100 µl elution buffer (25 mM TRIS-HCl, 300 mM NaCl, 300 mM imidazole, 10% vol. glycerol, 0.25 % vol. Tween 20, pH 7.8) were added to wash protein off the beads. The bead elution buffer mix was then incubated for one minute with occasional gentle vortexing. Afterward, the reaction tube was placed in the magnetic rack again to remove the eluted protein. This step was repeated for a second and third elution step. The buffer of the eluted protein was exchanged to TRIS buffered saline (TBS - 25mM TRIS, 72mM NaCl, 1mM $CaCl_2$ at pH 7.2) in 0.5 ml 40k Zeba spin columns distributed by ThermoFisher Scientific (Pierce Biotechnology, Rockford, IL, USA, # 87767) or 0.5 ml 50k Amicon Centrifugal Filters (Merck KGaA, Darmstadt, Germany, #UFC5050BK). Concentrations were determined photospectrometrically with a NanoDrop and aliquots were frozen in liquid nitrogen.

**Magnetic tweezers instrument**

Measurements were performed on a custom-built MT setup that has been described previously (*7, 8*). In the MT, molecules are tethered between a flow cell (FC; see next section) surface and superparamagentic beads (Dynabeads M-270 Streptavidin, Invitrogen, Life Technologies, Carlsbad, CA, USA). Mounted above the FC is a pair of permanent magnets (5×5×5 $mm^3$ each; W-05-N50-G, Supermagnete, Gottmadingen, Germany) in vertical configuration (34). The distance between magnets and FC is controlled by a DC-motor (M-126.PD2, Physik Instrumente PI GmbH & Co KG, Karlsruhe, Germany) and the FC is illuminated by an LED (69647, Lumitronix LED Technik GmbH, Germany). Using a 40x oil immersion objective (UPLFLN 40x, Olympus, Japan) and a CMOS sensor camera with 5120 x 5120 pixels (5120 x 5120 pixels, CP80-25-M-72, Optronis, Kehl, Germany) a field of view of approximately 680 × 680 $\mu m^2$ is imaged at a frame rate of 72 Hz. To control the focus and to create the look-up table required for tracking the bead positions in z, the objective is mounted on a piezo stage (Pifoc P-726.1CD, Physik Instrumente PI GmbH & Co KG, Karlsruhe, Germany). Images are read out with a frame grabber (microEnable 5 ironman VQ8-CXP6D, Silicon Software, Mannheim, Germany) and analyzed with an open-source tracking software (*9, 10*). The tracking accuracy of our setup is ≈ 1 nm in (x,y,z), as determined by tracking non-magnetic polystyrene beads, after baking them onto the flow cell surface. Force calibration was performed by analysis of the transverse fluctuations of long DNA tethers (*11*). Importantly, for the small extension changes on the length scales of our protein tethers, the force stays constant to very good approximation (to better than $10^{-4}$ relative change (*12*)). The largest source of force uncertainty is due to bead-to-bead variation, which is on the order of ≤ 10% for the beads used in this study (*13, 14*).

**Flowcell preparation and magnetic tweezers measurements**

Flowcells (FCs) were prepared as described previously (*12*). For the bottom slides, high precision microscope cover glasses (24 mm x 60 mm x 0.17 mm, Carl Roth) were amino-silanized for further functionalization (equal to AFM surface preparation). They were coated with sulfo-SMCC (*15*) (sulfosuccinimidyl 4-(N-maleimidomethyl)cyclohexane-1-carboxylate; sulfo-SMCC, ThermoFisher Scientific, Pierce Biotechnology, Rockford, IL, USA, # 22322). For this purpose, 180 µl sulfo-SMCC (10 mM in 50 mM Hepes buffer, pH 7.4) was applied to one amino-silanized slide that was sandwiched with another slide and incubated for 45 min. Unbound sulfo-SMCC was removed by rinsing with Milli-Q. Next, elastin-like polypeptide (ELP) linkers (*16*) with a sortase motif at their C-terminus were coupled to the maleimide of the sulfo-SMCC via a single cysteine at their N-terminus, by sandwiching two slides with 100 µl ELP linkers (in 50 mM Disodium phosphate buffer with 50mM NaCl and 10mM EDTA, pH 7.2) and incubating them for 60 min. Subsequently, after further Milli-Q rinsing to remove unbound ELP linkers, free sulfo-SMCC was neutralized with free cysteine (10 mM in 50 mM disodium phosphate buffer with 50 mM NaCl and 10 mM EDTA, pH 7.3). 1 µm diameter polystyrene beads dissolved in ethanol were applied to the glass slides. After the ethanol evaporated, beads were baked onto the glass surface for 5 min at ≈ 80°C to serve as reference beads during the measurement. FCs were assembled from an ELP-functionalized bottom slide and an unfunctionalized high-precision microscope cover

glass slide with two holes (inlet and outlet) on either side serving as top slide. Both slides were separated by a layer of parafilm (Pechiney Plastic Packaging Inc., Chicago, IL, USA), which was cut out to form a 50 µl channel. FCs were incubated with 1% (v/v) casein solution (# C4765-10ML, Sigma-Aldrich) for 2 h and flushed with 1 ml buffer (25 mM TRIS, 72 mM NaCl, 1 mM CaCl2, pH 7.2 at room temperature).
CoA-biotin (# S9351 discontinued, New England Biolabs, Frankfurt am Main, Germany) was coupled to the ybbR-tag at the C-terminus of the fusion protein constructs in a 90 - 120 min bulk reaction in the presence of 4 µM sfp phosphopantetheinyl transferase (*17*) and 100 mM $MgCl_2$ at room temperature (≈ 22°C). Proteins were diluted to a final concentration of about 50 nM in 25 mM TRIS, 72 mM NaCl, 1 mM $CaCl_2$, pH 7.2 at RT. To couple the N-terminus of the fusion proteins carrying three glycines to the C-terminal LPETGG motif of the ELP-linkers, 100 µl of the protein mix was flushed into the FC and incubated for 24 min in the presence of 1.3 µM evolved pentamutant sortase A from Staphylococcus aureus (*18, 19*). Unbound proteins were flushed out with 1 ml measurement buffer (25 mM TRIS, 72 mM NaCl, 1 mM CaCl2, 0.1% (v/v) Tween-20, pH 7.2). Finally, commercially available streptavidin-coated paramagnetic beads (Dynabeads™ M-270 Streptavidin, Invitrogen, Life Technologies, Carlsbad, CA, USA) were added into the FC and incubated for 30 s before flushing out unbound beads with 1 ml measurement buffer. Receptor-ligand binding and unbinding under force was systematically investigated by subjecting the protein tethers to (2 -30) min long plateaus of constant force, which was gradually increased in steps of 0.2 or 0.3 pN. All measurements were conducted at room temperature.

**Data analysis of MT traces**

MT traces were selected on the basis of the characteristic ACE2 two-step unfolding pattern above 25 pN, conducted at the end of each experiment. For each trace, (x,y)-fluctuations were also checked to avoid inclusion of tethers that exhibit inter-bead or bead-surface interactions, which would also cause changes in x or y. Non-magnetic reference beads were tracked simultaneously with magnetic beads and reference traces were subtracted for all measurements to correct for drift. Extension time traces were smoothed to one second with a moving average filter to reduce noise. All analyses were performed with custom scripts in MATLAB.

**Molecular dynamics simulations**

To provide a complementary microscopic view of the RBD:ACE2 complex, we carried out molecular dynamics (MD) simulations employing NAMD 3 (*20*). Simulations were prepared using VMD (*21*) and its QwikMD (*22*) interface. The structure of the complexes were prepared following established protocols (*23*). As a starting point, we used the crystallographic structure of the wt SARS-CoV-2 RBD:ACE2 complex from the protein data bank (PDB ID: 6m0j) (*24*). The structure for the VOCs Alpha, Beta, Gamma, Delta and Omicron were obtained using Modeller (*25*), with standard parameters and implementing the mutations described at for VOCs Alpha, Beta, Gamma and Delta.

Omicron VOC model was constructed according to the following mutations found on RBD: G339D, S371L, S373P, S375F, K417N, N440K, G446S, S477N, T478K, E484A, Q493R, G496S, Q498R, N501Y, Y505H.

Employing advanced run options of QwikMD, structural models were solvated and the net charge of the proteins were neutralized using a 75 mM salt concentration of sodium chloride, which were randomly arranged in the solvent. All simulations were performed employing the NAMD MD package (*20*) and run on NVIDIA DGX-A100-based cluster nodes at Auburn University. The CHARMM force field (*26, 27*) along with the TIP3 water model (*28*) was used for all systems. The simulations were performed with periodic boundary conditions in the NpT ensemble with temperature maintained at 300 K and pressure at 1 bar using Langevin dynamics. A distance cut-off of 12.0 Å was applied to short-range, non-bonded interactions, whereas long-range electrostatic interactions were treated using the particle-mesh Ewald (PME) (*29*) method. The equations of motion were integrated using the r-RESPA multiple time step scheme (*30*) to update the van der Waals interactions every two steps and electrostatic interactions every four steps. The time step of integration was chosen to be 4 fs for all production simulations performed, and 2 fs for all equilibration runs. For the 4 fs simulations, hydrogen mass repartitioning was done using psfgen in VMD. Before the MD simulations all the systems were submitted to an energy minimization protocol for 5,000 steps.

MD simulations with position restraints in the protein backbone atoms were performed for 1.0 ns and served to pre-equilibrate systems before the 10 ns equilibrium MD runs, which served to evaluate structural model stability. During the 1.0 ns pre-equilibration the initial temperature was set to zero and was constantly increased by 1 K every 1,000 MD steps until the desired temperature (300 K) was reached. Production runs with no restraints on the system were performed for 200 ns, in five replicas, for each system, totaling 1 microsecond per system.

**Analysis of molecular dynamics simulation**

Analyses of MD trajectories were carried out using Dynamical Network Analysis (*31*), using custom python code and VMD (*21*) and its plug-ins. In Dynamical Network Analysis, (*31*) a network is defined as a set of , and each node represents an amino acid residue. Each node's position is given by the residue's α-carbon. Edges connect pairs of nodes if their corresponding residues are in contact and 2 non-consecutive residues are said to be in contact if they are within 4.5 Å of each other for at least 75% of (*31*) analyzed frames. To ensure a broad sampling of our systems, each of the five 200-ns MD trajectories for each system were split in 5 ns windows, and only the last 15 windows, or 75 ns, were used for analysis. Moreover, for the analysis of total correlation in RBD:ACE2 interfaces, we filtered out all contacts that presented presented average correlation of motion smaller than 0.2 in order to reduce noise and remove weak transient interactions from the analysis.

**Bootstrapping for confidence intervals and significance testing**

The confidence intervals (CIs) for the mean total correlation reported in Figure 3D were determined using bootstrapping and the bias-corrected accelerated method (*32*), as implemented in SciPy (*33*). The same method was used to determine the CIs presented in Supplementary Figures S3 and S10, all CIs at a 90% confidence level.

To test if the total correlation distributions from distinct VOCs had significantly different means (Supplementary Figure S10A), we used the non-parametric bootstrapping technique for hypothesis testing, with 10,000 samplings, as proposed by Efron, *et al.* (*32*). The same technique was used on the experimental data for $F_{1/2}$ (Supplementary Figure S10B) and we obtained similar results as with the parametric *t*-test.

**Two-dimensional network of protein interface**

The two-dimensional representation in Supplementary Figure S7 were created by first mapping the 3D positions of residues of the RBD near the RBD:ACE2 interface in the WT system to a 2D space. This was done using a principal component analysis transformation as implemented in Scikit-Learn (*34*). The plots were created using the interface between NetworkX (*35*) and matplotlib (*36*) in Python. Mean correlations were calculated as described above, and confidence intervals for S7A were calculated using the bias-corrected accelerated method for bootstrapping.

**FULL PROTEIN SEQUENCES**

Sortase N-Tag
ACE2
85 aa linker
His6-Tag
ybbR
Avitag
Fgɣ

**Basis construct:**
**pT7CFE1-MGGG-ACE2-85aa-linker-[INSERT]-HIS-ybbr-AviTag-Fgɣ**

MGGGSSSTIEEQAKTFLDKFNHEAEDLFYQSSLASWNYNTNITEENVQNMNNAGDKWSAFLKEQS
TLAQMYPLQEIQNLTVKLQLQALQQNGSSVLSEDKSKRLNTILNTMSTIYSTGKVCNPDNPQECL
LLEPGLNEIMANSLDYNERLWAWESWRSEVGKQLRPLYEEYVVLKNEMARANHYEDYGDYWRGDY
EVNGVDGYDYSRGQLIEDVEHTFEEIKPLYEHLHAYVRAKLMNAYPSYISPIGCLPAHLLGDMWG
RFWTNLYSLTVPFGQKPNIDVTDAMVDQAWDAQRIFKEAEKFFVSVGLPNMTQGFWENSMLTDPG
NVQKAVCHPTAWDLGKGDFRILMCTKVTMDDFLTAHHEMGHIQYDMAYAAQPFLLRNGANEGFHE
AVGEIMSLSAATPKHLKSIGLLSPDFQEDNETEINFLLKQALTIVGTLPFTYMLEKWRWMVFKGE
IPKDQWMKKWWEMKREIVGVVEPVPHDETYCDPASLFHVSNDYSFIRYYTRTLYQFQFQEALCQA
AKHEGPLHKCDISNSTEAGQKLFNMLRLGKSEPWTLALENVVGAKNMNVRPLLNYFEPLFTWLKD
QNKNSFVGWSTDWSPYADGATSGGGGSAGGSGSGSSGGSSGASGTGTAGGTGSGSGTGSGGGSGG
GSEGGGSEGGGSEGGGSEGGGSEGGGSGGGSESGGSSA**[INSERT]**SGHHHHHHTDSLEFIASKL
AASGLNDIFEAQKIEWHEGSGEGQQHHLGGAKQAGDV*

**Inserts:**

**SARS-CoV-1-RBD**

RVVPSGDVVRFPNITNLCPFGEVFNATKFPSVYAWERKKISNCVADYSVLYNSTFFSTFKCYGVS
ATKLNDLCFSNVYADSFVVKGDDVRQIAPGQTGVIADYNYKLPDDFMGCVLAWNTRNIDATSTGN
YNYKYRYLRHGKLRPFERDISNVPFSPDGKPCTPPALNCYWPLNDYGFYTTTGIGYQPYRVVVLS
FELLNAPATVCGPKLSTDLIKNQCVNF

**SARS-CoV-2-RBD**

SNFRVQPTESIVRFPNITNLCPFGEVFNATRFASVYAWNRKRISNCVADYSVLYNSASFSTFKCY
GVSPTKLNDLCFTNVYADSFVIRGDEVRQIAPGQTGKIADYNYKLPDDFTGCVIAWNSNNLDSKV
GGNYNYLYRLFRKSNLKPFERDISTEIYQAGSTPCNGVEGFNCYFPLQSYGFQPTNGVGYQPYRV
VVLSFELLHAPATVCGPKKSTNLVKN

**Alpha-RBD**

SNFRVQPTESIVRFPNITNLCPFGEVFNATRFASVYAWNRKRISNCVADYSVLYNSASFSTFKCY
GVSPTKLNDLCFTNVYADSFVIRGDEVRQIAPGQTGKIADYNYKLPDDFTGCVIAWNSNNLDSKV
GGNYNYLYRLFRKSNLKPFERDISTEIYQAGSTPCNGVEGFNCYFPLQSYGFQPTYGVGYQPYRV
VVLSFELLHAPATVCGPKKSTNLVKN

**Beta-RBD**

SNFRVQPTESIVRFPNITNLCPFGEVFNATRFASVYAWNRKRISNCVADYSVLYNSASFSTFKCY
GVSPTKLNDLCFTNVYADSFVIRGDEVRQIAPGQTGNIADYNYKLPDDFTGCVIAWNSNNLDSKV
GGNYNYLYRLFRKSNLKPFERDISTEIYQAGSTPCNGVKGFNCYFPLQSYGFQPTYGVGYQPYRV
VVLSFELLHAPATVCGPKKSTNLVKN

**Gamma-RBD**

SNFRVQPTESIVRFPNITNLCPFGEVFNATRFASVYAWNRKRISNCVADYSVLYNSASFSTFKCY
GVSPTKLNDLCFTNVYADSFVIRGDEVRQIAPGQTGTIADYNYKLPDDFTGCVIAWNSNNLDSKV
GGNYNYLYRLFRKSNLKPFERDISTEIYQAGSTPCNGVKGFNCYFPLQSYGFQPTYGVGYQPYRV
VVLSFELLHAPATVCGPKKSTNLVKN

**Delta-RBD**

SNFRVQPTESIVRFPNITNLCPFGEVFNATRFASVYAWNRKRISNCVADYSVLYNSASFSTFKCY
GVSPTKLNDLCFTNVYADSFVIRGDEVRQIAPGQTGKIADYNYKLPDDFTGCVIAWNSNNLDSKV
GGNYNYRYRLFRKSNLKPFERDISTEIYQAGSKPCNGVEGFNCYFPLQSYGFQPTNGVGYQPYRV
VVLSFELLHAPATVCGPKKSTNLVKN

| | wt (G614D) | Alpha | Beta | Gamma | Delta |
|---|---|---|---|---|---|
| $F_{1/2,\ fit}$ ± std (pN) | 3.8 ± 0.4 | 4.5 ± 0.3 | 3.8 ± 0.3 | 4.0 ± 0.6 | 3.8 ± 0.3 |
| $\Delta z$ ± std (nm) | 10.4 ± 3.6 | 14 ± 6.4 | 11.4 ± 1.8 | 13.2 ± 5.3 | 12.5 ± 3.0 |
| $\tau_{0,diss}$ ± std (s) | 0.06 ± 0.17 | 0.01 ± 0.02 | 0.02 ± 0.06 | 0.02 ± 0.04 | 0.01 ± 0.02 |
| $\tau_{0,bound}$ ± std (s) | 121 ± 284 | 464 ± 740 | 218 ± 315 | 552 ± 1178 | 427 ± 707 |
| Kd | $532 \cdot 10^{-6}$ | $30 \cdot 10^{-6}$ | $110 \cdot 10^{-6}$ | $31 \cdot 10^{-6}$ | $27 \cdot 10^{-6}$ |
| $F_{1/2,\ dwelltimes}$ ± std (pN) | 3.8 ± 0.5 | 4.4 ± 0.4 | 3.9 ± 0.3 | 4.0 ± 0.5 | 3.8 ± 0.2 |

**Supplementary Table T1. Fit parameters from equilibrium and kinetic interaction analysis of SARS-CoV-2 VOCs with ACE2 in MT.** $F_{1/2}$, fit and $\Delta z$ are the mean fit parameters of all molecules from fitting equation 1 to the force-dependent fraction in the dissociated state. $\tau_{0,diss}$ and $\tau_{0,bound}$ are the mean lifetimes at zero force of all molecules, determined by fitting equations 2 and 3 to the force-dependent dwell times. Kd is the resulting dissociation constant determined as $\tau_{0,diss}/\tau_{0,bound}$. $F_{1/2}$, dwell times is the mean midpoint force determined as the intersection of the force-dependent dwell times in the bound and dissociated state for individual molecules.

| Study | | SARS-CoV-1 | SARS-CoV-2 | Alpha | Beta | Gamma | Delta | Method and Comments |
|---|---|---|---|---|---|---|---|---|
| Lan et al. (*24*) | ⤙ | $K_d$ = 31 nM<br>$k_{sol,off}$ = 4.3×10$^{-2}$ s$^{-1}$<br>$k_{sol,on}$ = 1.4×10$^{6}$ s$^{-1}$M$^{-1}$ | $K_d$ = 4.7 nM<br>$k_{sol,off}$ = 6.5×10$^{-3}$ s$^{-1}$<br>$k_{sol,on}$ = 1.4×10$^{6}$ s$^{-1}$M$^{-1}$ | | | | | Surface-plasmon resonance, immobilized human ACE2 (residues Ser19–Asp615) with the SARS-CoV-2 RBD (D (residues Arg319–Phe541)) and SARS-CoV RBD |
| Shang et al. (*37*) | ◀ | $K_d$ =185 nM<br>$k_{sol,off}$ = 3.37×10$^{-2}$ s$^{-1}$<br>$k_{sol,on}$ = 2.01×10$^{5}$ s$^{-1}$M$^{-1}$ | $K_d$ =44.2 nM<br>$k_{sol,off}$ = 7.75×10$^{-3}$ s$^{-1}$<br>$k_{sol,on}$ = 1.75×10$^{5}$ s$^{-1}$M$^{-1}$ | | | | | Surface-plasmon resonance, purified recombinant RBDs were covalently immobilized on the sensor chip through their amine groups and purified recombinant ACE2 (residues 1–615) flowed over the RBDs |
| Starr et al. (*38*) | ▶ | $K_d$ =575 nM | $K_d$ =92 nM | | | | | Biolayer interferometry binding analysis between RBDs of SARS-CoV-2 |

| | | | | | | | | |
|---|---|---|---|---|---|---|---|---|
| | | | | | | | | (328-531) or SARS-CoV-1 (306-575) and ACE2 (residues 1–615) |
| Walls et al. (*39*) | ⬟ | $K_d$ = 5.0 ± 0.1 nM<br>$k_{sol,off}$ = (8.7 ± 5.1) $\times 10^{-4}$ $s^{-1}$<br>$k_{sol,on}$ = (1.7 ± 0.7) $\times 10^{5}$ $s^{-1}M^{-1}$ | $K_d$ = 1.2 ± 0.1 nM<br>$k_{sol,off}$ = (1.7 ± 0.8) $\times 10^{-4}$ $s^{-1}$<br>$k_{sol,on}$ = (2.3 ± 1.4) $\times 10^{5}$ $s^{-1}M^{-1}$ | | | | | Biolayer interferometry binding analysis of the hACE2 ectodomain (residues 1–614) to immobilized SARS-CoV-2 (N-terminal 328RFPN331 and C-terminal 530STNL533) or SARS-CoV RBD (306RVVPSG311 and C-terminal 571LDISP575) |
| Wang et al. (*40*) | Y | $K_d$ = 408.7 ± 11 nM<br>$k_{sol,off}$ = (1.9 ± 0.4) $\times 10^{-3}$ $s^{-1}$<br>$k_{sol,on}$ = (2.9 ± 0.2) $\times 10^{5}$ $s^{-1}M^{-1}$ | $K_d$ = 94.6 ± 7 nM<br>$k_{sol,off}$ = (3.8 ± 0.2) $\times 10^{-3}$ $s^{-1}$<br>$k_{sol,on}$ = (4.0 ± 0.2)$\times 10^{4}$ $s^{-1}M^{-1}$ | | | | | Surface-plasmon resonance; assessing binding affinity of SARS-CoV-2 RBD (319−541) or SARS-RBD (residues 306-527, GenBank: NC_004718) to ACE2 (residues 19−615) |

| | | | | | | | | |
|---|---|---|---|---|---|---|---|---|
| Wrapp et al. (*41*) | \| | $K_d$ = 325.8 nM<br>$k_{sol,off}$ = 0.112×10$^{-3}$ s$^{-1}$<br>$k_{sol,on}$ = 3.62×10$^{5}$ s$^{-1}$M$^{-1}$ | $K_d$ = 34.6 nM<br>$k_{sol,off}$ = 4.7 ×10$^{-3}$ s$^{-1}$<br>$k_{sol,on}$ = 1.36 ×10$^{5}$ s$^{-1}$M$^{-1}$ | | | | | Surface-plasmon resonance; assessing binding of ACE2 (residues 1−615) to the SARS-CoV-2 RBD (319−591) and SARS-CoV RBD (306−577) |
| Barton et al. (*42*) | — | | Calc: 74.4 nM<br>$k_{sol,off}$ = 0.0668 s$^{-1}$<br>$k_{sol,on}$ = 0.90 µM$^{-1}$s$^{-1}$<br><br>Measured: 62.6 nM | 7.0 nM<br>$k_{sol,off}$ = 0.0111 s$^{-1}$<br>$k_{sol,on}$ = 1.59 µM$^{-1}$s$^{-1}$<br><br>Measured: 5.5 nM | 20.0 nM<br>$k_{sol,off}$ = 0.0291 s$^{-1}$<br>$k_{sol,on}$ = 1.46 µM$^{-1}$s$^{-1}$<br><br>Measured: 17.4 nM | 13.5 nM<br>$k_{sol,off}$ = 0.0211 s$^{-1}$<br>$k_{sol,on}$ = 1.56 µM$^{-1}$s$^{-1}$<br><br>Measured: 12.2 nM | | Surface-plasmon resonance; KD between SARS-CoV-2 RBDs (328−550) and ACE2 (residues 1−615) calculated from on and off rate and measured |
| Laffeber et al. (*43*) | ★ | | 17 nM<br>$k_{sol,off}$ = 7.8 ×10$^{-3}$ s$^{-1}$<br>$k_{sol,on}$ = 4.5 ×10$^{5}$ M$^{-1}$s$^{-1}$ | 2.4 nM<br>$k_{sol,off}$ = 1.3 ×10$^{-3}$ s$^{-1}$<br>$k_{sol,on}$ = 5.7 ×10$^{5}$ M$^{-1}$s$^{-1}$ | 5.8 nM<br>$k_{sol,off}$ = 4.3 ×10$^{-3}$ s$^{-1}$<br>$k_{sol,on}$ = 7.6 ×10$^{5}$ M$^{-1}$s$^{-1}$ | | | Surface-plasmon resonance; KD between RBD (reference genome Wuhan-HU-1; residues 333–529) and ACE2 (residues 18–615) |

| | | | | | | | | |
|---|---|---|---|---|---|---|---|---|
| Gong et al. (*44*) | ≻ | | *161 nM*<br>$k_{sol,off} = 6.88 \times 10^{-3}\ s^{-1}$<br>$k_{sol,on} = 4.27 \times 10^{4}\ M^{-1}s^{-1}$ | Flow: wt/5.43<br><br>BLI:<br>41.5 nM<br>$k_{sol,off} = 1.49 \times 10^{-3}\ s^{-1}$<br>$k_{sol,on} = 3.59 \times 10^{4}\ M^{-1}s^{-1}$ | Flow: wt/3.56 | Flow: wt/ 4.24 | Flow: wt/ 2.34 | Biolayer interferometry (BLI) for wt and Alpha (both residues 319–541) and ACE2 (1−615); binding of ACE2 to cell-surface Spike by flow cytometry was measured for Alpha, Beta, Gamma and Delta |
| Rajah et al. (*45*) | + | | 14.05 ug/ml | 0.44 ug/ml | | | 1.64 ug/ml | EC50 for binding of full ACE2 on cell-surface to SARS-CoV-2 Spike (GenBank: QHD43416.1) by flow cytometry |
| Gobeil et al. (*46*) | × | | 218.29 nM<br>Kon = 6.23 *10^4<br>koff = 1.36 *10^-2 | 51.8 nM<br>Kon = 5.48 *10^4,<br>koff = 2.84*10^-3 | 93.72 nM<br>Kon = 9.54 *10^4<br>koff = 8.96 *10^-3 | | | Surface-plasmon resonance, spike (1 to 1208, GenBank MN908947) binding to the ACE2 receptor ectodomain |

| | | | | | | | | |
|---|---|---|---|---|---|---|---|---|
| Ren et al. (*47*) | ● | | 5.5 nM<br>$k_{sol,off}$ = 8.525 $\times 10^{-3}$ $s^{-1}$<br>$k_{\mathrm{sol,on}}$ = 1.550 $\times 10^{6}$ $M^{-1}s^{-1}$ | | | | 2.67 nM<br>$k_{sol,off}$ = 4.606 $\times 10^{-3}$ $s^{-1}$<br>$k_{\mathrm{sol,on}}$ = 1.767 $\times 10^{6}$ $M^{-1}s^{-1}$ | Surface-plasmon resonance; KD between WT or Delta SARS-CoV-2 RBD (residues Arg319–Phe541) and the N-terminal peptidase domain of human or mouse ACE-2 (residues Ser19–Asp615) |
| McCallum et al. (*48*) | ⅄ | | SPR: 78 nM<br>$k_{sol,off}$ = 6.7 $\times 10^{-3}$ $s^{-1}$<br>$k_{\mathrm{sol,on}}$ = 7.7 $\times 10^{4}$ $M^{-1}s^{-1}$<br><br>BLI: 147 nM<br>$k_{sol,off}$ = 1.4 $\times 10^{-2}$ $s^{-1}$<br>$k_{\mathrm{sol,on}}$ = 9.5 $\times 10^{4}$ $M^{-1}s^{-1}$ | SPR: 15 nM<br>$k_{sol,off}$ = 1.2 $\times 10^{-3}$ $s^{-1}$<br>$k_{\mathrm{sol,on}}$ = 7.5 $\times 10^{4}$ $M^{-1}s^{-1}$<br><br>BLI: 26 nM<br>$k_{sol,off}$ = 2.8 $\times 10^{-3}$ $s^{-1}$<br>$k_{\mathrm{sol,on}}$ = 1.3 $\times 10^{5}$ $M^{-1}s^{-1}$ | | | SPR: 63 nM<br>$k_{sol,off}$ = 4.3 $\times 10^{-3}$ $s^{-1}$<br>$k_{\mathrm{sol,on}}$ = 5.9 $\times 10^{4}$ $M^{-1}s^{-1}$<br><br>BLI: 180 nM<br>$k_{sol,off}$ = 1.1 $\times 10^{-2}$ $s^{-1}$<br>$k_{\mathrm{sol,on}}$ = 7.4 $\times 10^{4}$ $M^{-1}s^{-1}$ | Surface-plasmon resonance and BLI; KD between RBD (N-328-RFPN-331 and 528-KKST-531) and ACE2 (residues 1-615) |

**Supplementary Table T2. Equilibrium binding data for SARS-CoV-1 or SARS-CoV-2 RBD and variants of concerns binding to ACE2 shown in Figure 2D.**

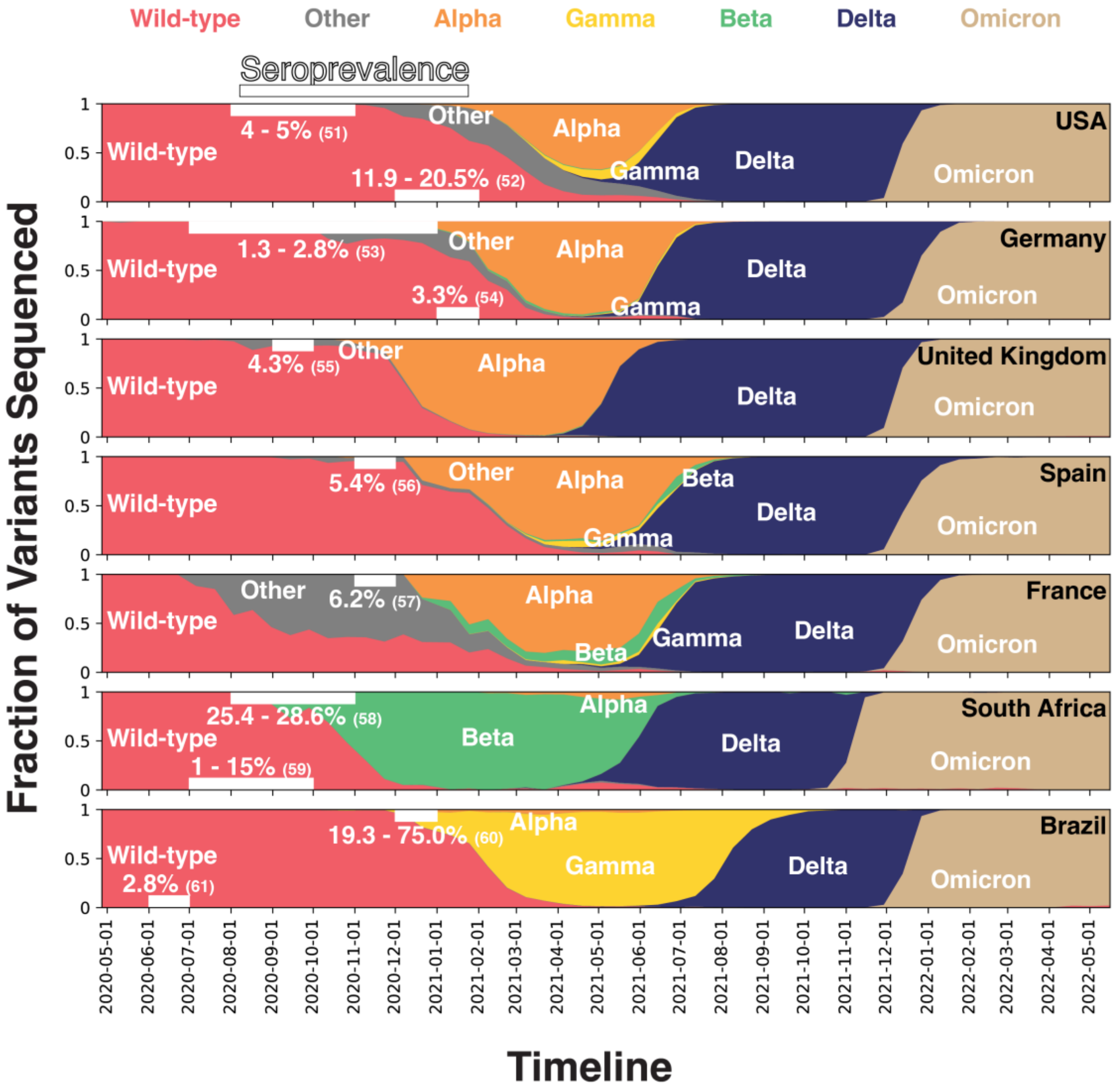


**Supplementary Fig. S1.** Fraction of variants of concern sequenced, provided by GISAID (*49*) via CoVariants.org (*50*), in seven representative countries over time between May 2020 and May 2022. The data provide a proxy to determine which VOCs become dominant in the respective countries over time. In white the nationwide seroprevalence determined by selected studies is shown to indicate the seroprevalence of the general public in the marked timespan (*51-62*). The timespan of the seroprevalence studies was chosen at the period of the transition between the initial wild type strain losing dominance over the upcoming VOCs. In Germany, the United Kingdom, Spain, France, and the United States, Alpha replaced the wild type and became the dominantly circulating and sequenced variant. This was at a time where no widespread population immunity prevailed, suggesting that Alpha has a significant advantage in largely immunologically naïve populations. This might imply that higher force stability contributes a fitness advantage for Alpha, in an immunologically naïve population. In conclusion this could hint at force stability as a possible selection factor for VOCs. In contrast Beta or Gamma, as seen in South Africa and Brazil, only became dominant in regions with higher population immunity, a setting where amino acid substitutions contributing to immune evasion might provide a more important fitness advantage compared to higher force stability.

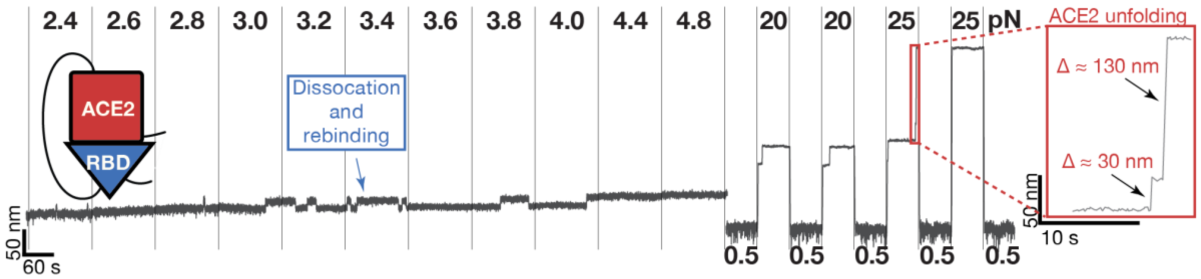


**Supplementary Fig. S2. Unfolding of the tethered ligand constructs provides a molecular fingerprint pattern.** Extension-time trace of an RBD:ACE2 tethered ligand construct in MT at different forces, indicated at the top of the trace. MT traces are selected for analysis based on dissociation and rebinding pattern visible at forces < 5 pN and specific, irreversible two-step ACE2 unfolding patterns between 20 and 25 pN, shown in detail in the inset. The ACE2 unfolding pattern was previously identified and assigned using magnetic tweezers and AFM force spectroscopy (*1*).

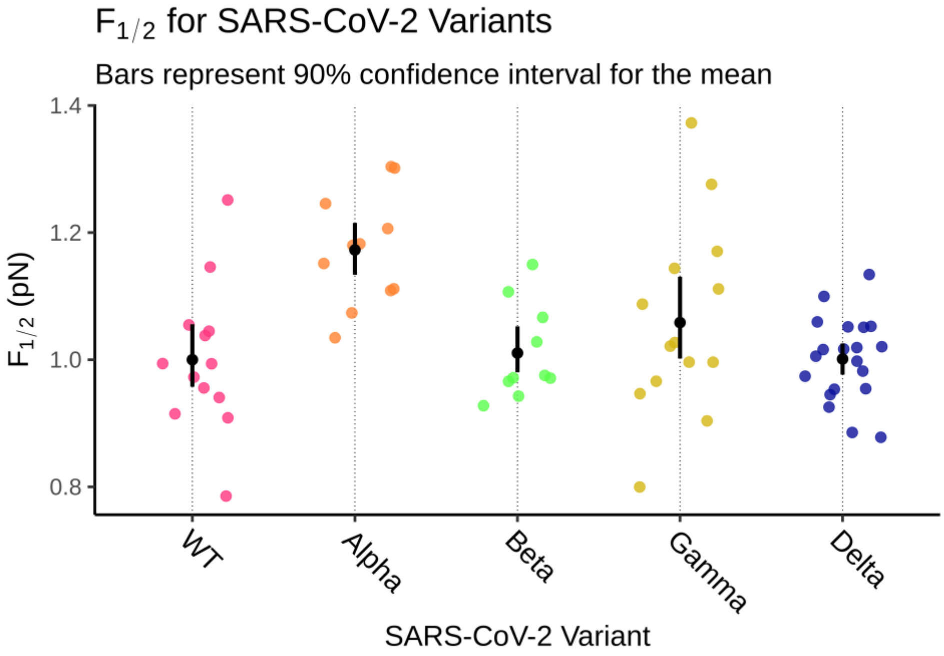


**Supplementary Fig. S3.** Small colored circles are values of midpoint forces determined for wt, Alpha, Beta, Gamma, and Delta VOCs (Figure 2B), normalized by the mean wt $F_{1/2}$. Black dots and errors bars are the mean and confidence intervals for $F_{1/2}$ estimated with the same bootstrapping approach used for total correlations (Figure 3D and Supplementary Materials and Methods). The Alpha VOC is significantly different from all other VOCs and wt (see Figure S10 for p-values and SI methods for details).

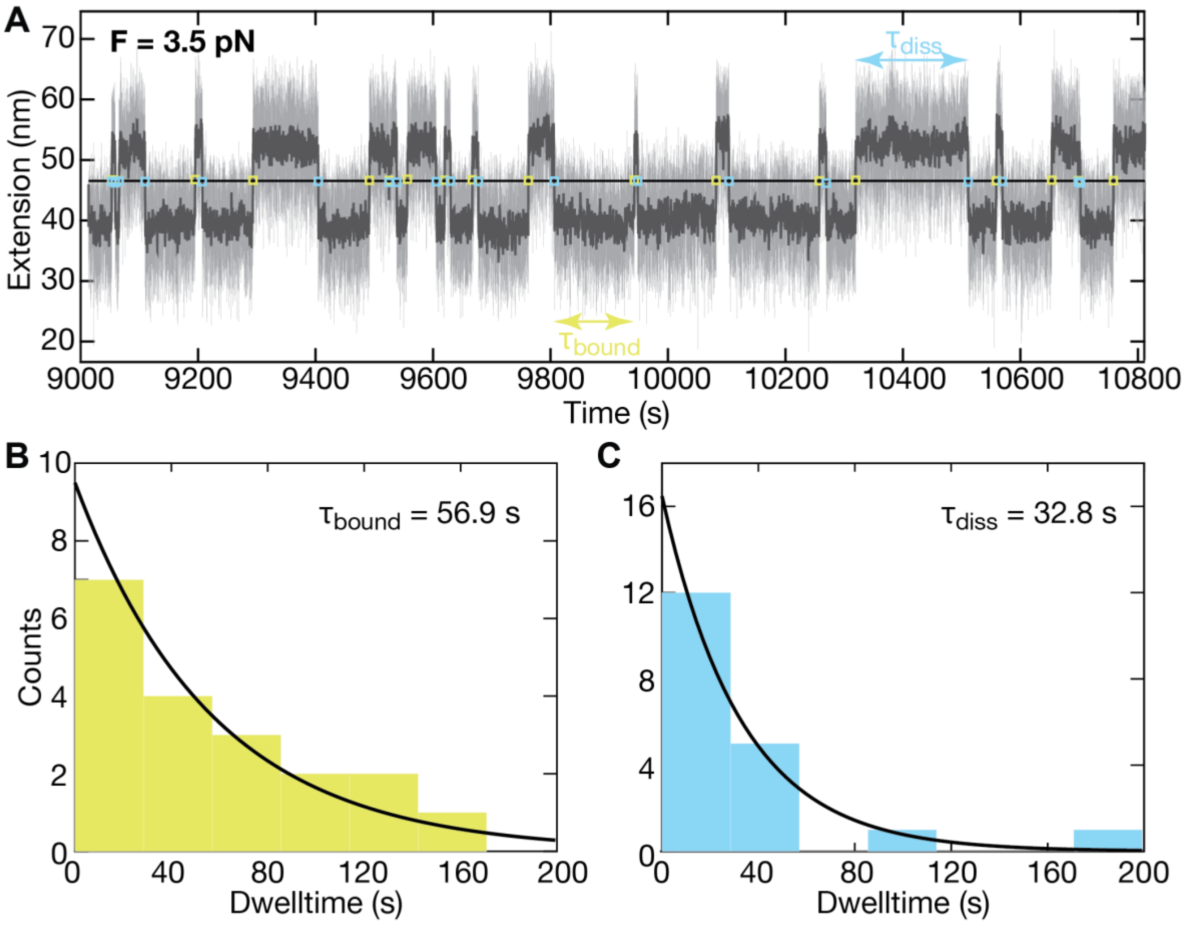


**Supplementary Fig. S4. Dwell time analysis of the tethered ligand extension time traces in MT. A** Short segment of an extension time trace measured for a SARS-CoV-2 RBD:ACE2 tethered ligand construct at a force of 3.5 pN. Raw data at 72 Hz are shown in light grey and filtered data (50 frame moving average) are shown in dark grey. Assignment of the dwell times is based on the filtered data. The black horizontal line is the threshold; green squares indicate the first data point after crossing the threshold from below, i.e. transition from the bound to the dissociated state; blue squares indicate the first data point after crossing the threshold from above, i.e. transition from the dissociated to the bound state. **B,C** Histograms of dwell times in the bound state (**B**) and dissociated state (**C**) obtained from the analysis shown in panels **A**. The dwell times are well described by single exponential fits, shown as solid lines. Insets show the mean dwell times from maximum likelihood fits of the single exponentials.

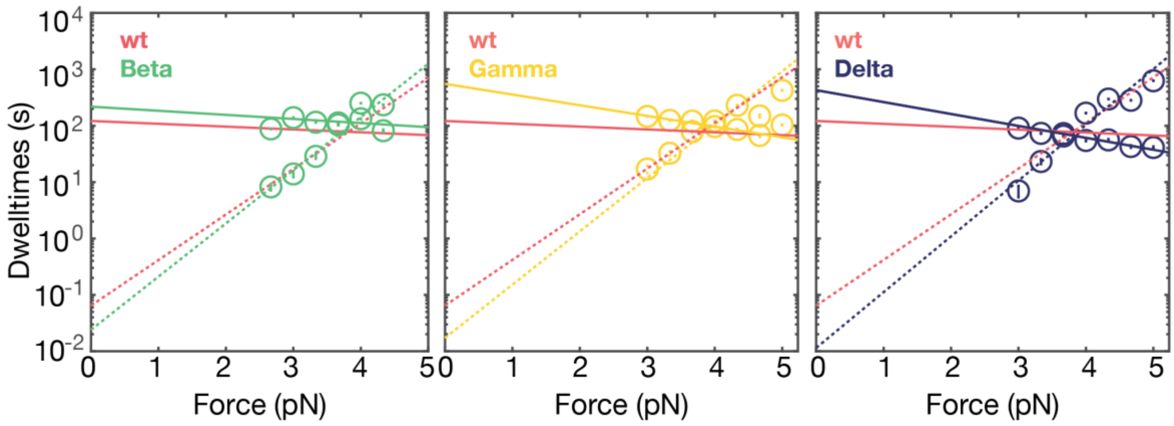


**Supplementary Fig. S5. Force-dependent dwelltimes for VOC and comparison to wt.** Mean life times at different levels of applied force in the dissociated and bound conformation of the RBD:ACE2 construct determined experimentally (circles) and exponential fits (solid line for the bound state lifetimes and dashed lines for the dissociated state lifetimes). For comparison, the fit lines for the wt are co-plotted in red. Even though mid forces are similar for all variants, slopes differ and result in different extrapolated life times at zero force.

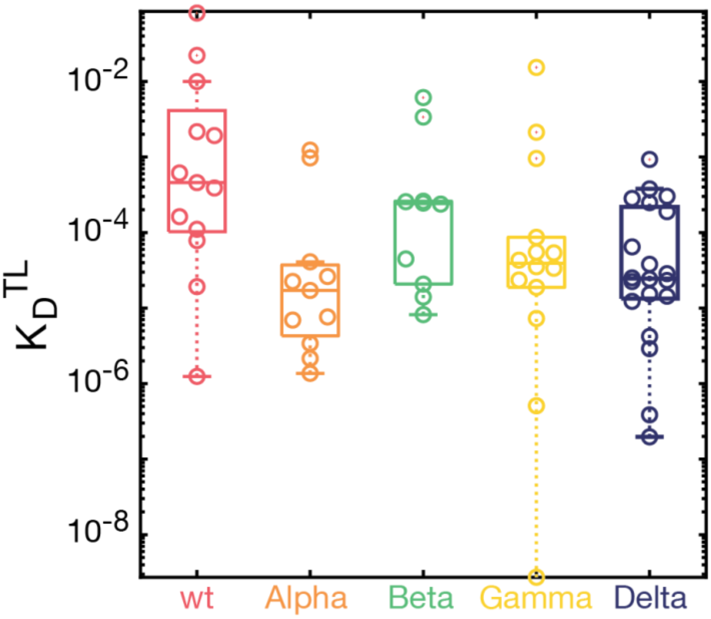


**Supplementary Fig. S6. $K_D^{TL}$ values determined for the tethered ligand constructs.** Tethered ligand dissociation constants $K_D^{TL}$ are obtained as $\tau_{0,diss}/\tau_{0,bound}$, i.e. as the ratio of the mean life times in the dissociated and bound states extrapolated to zero force. Small circles are values obtained for individual molecules. Boxes indicate the median and the the 25th and 75th percentile of the data. All VOC have lower $K_D$ values and, therefore, higher affinities for ACE2 than the wt.

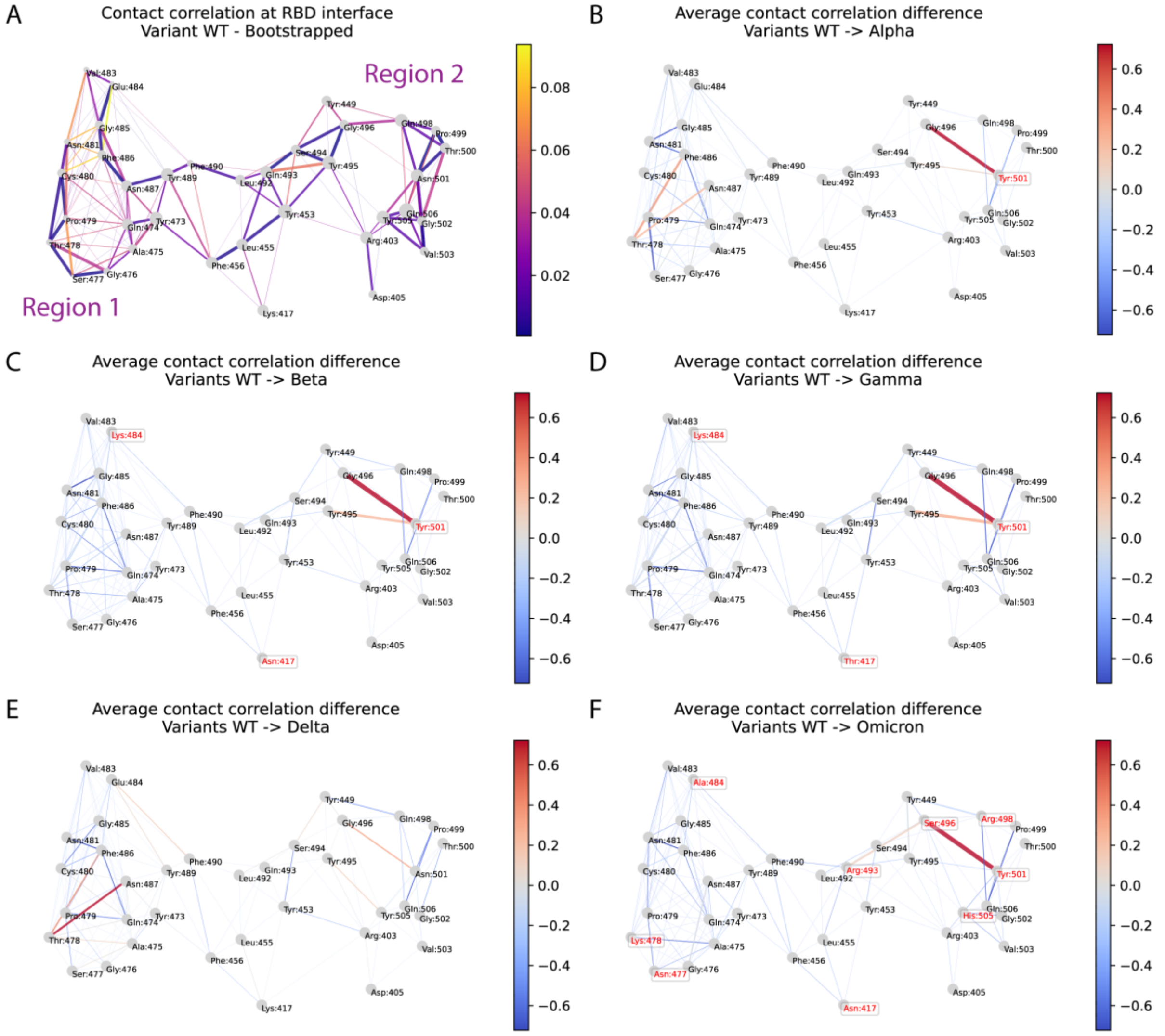


**Supplementary Fig. S7. Correlation difference maps for RBD interface.** The panels show 2D representations of all residues on the RBD surface that compose its interface with ACE2. **A** Edges indicate residues in contact. Edge thickness indicates the mean correlation between the motion of neighboring residues and is a proxy for the strength of their interactions. Edge color indicates the width of the confidence interval (CI) for the mean correlation, where stable interaction leads to narrower CIs (in purple) while irregular interaction leads to wider CIs (in yellow). Regions 1 and 2 are defined in Figure 3B. **B** through **F** Edge thickness and color indicate the difference between average correlation measurements for the WT system and for VOCs. Color scale varies from loss of correlation in blue to gain of correlation of motion in red.

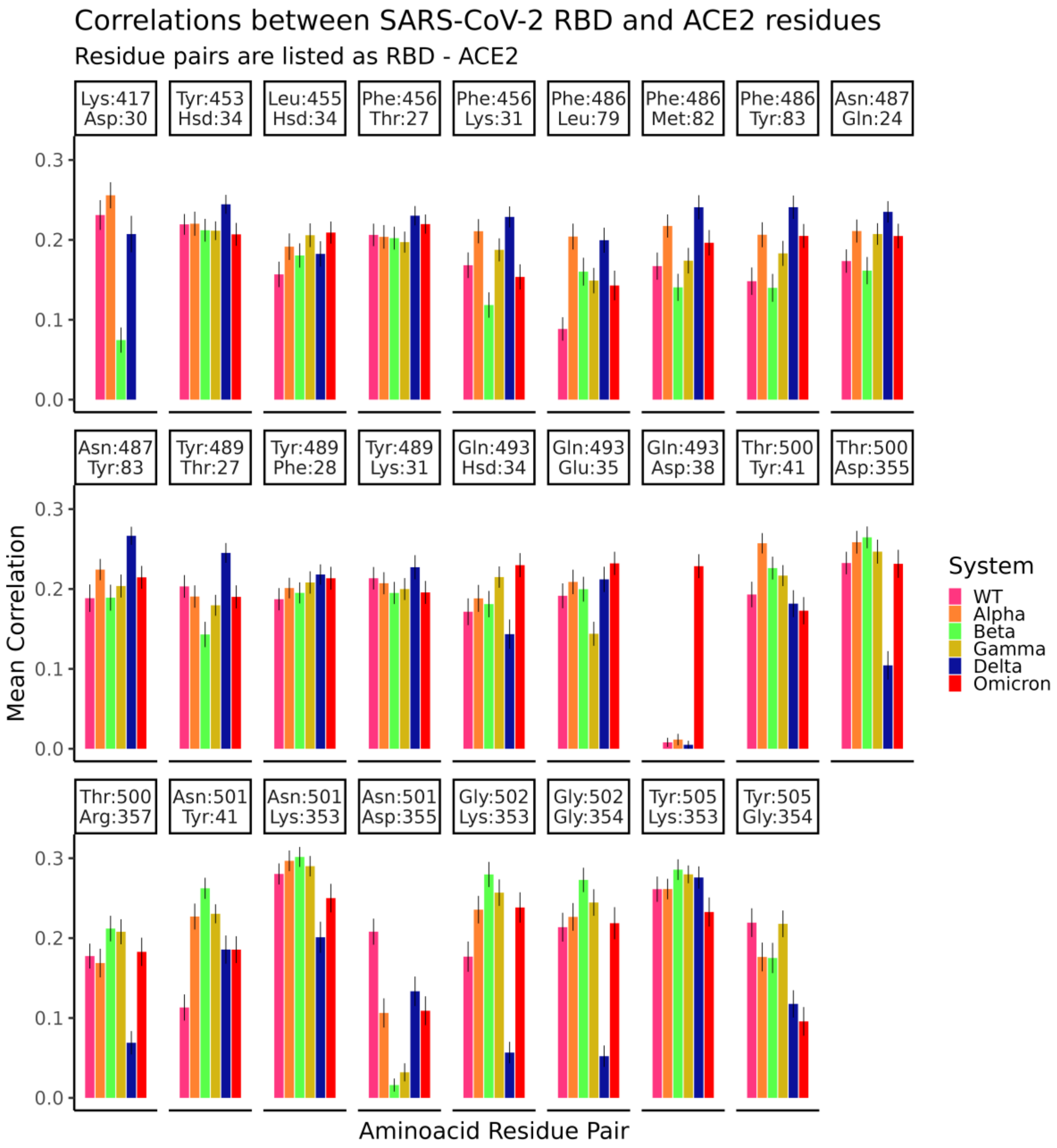


**Supplementary Fig. S8.** Generalized correlation between the motions of residue pairs across the RBD:ACE2 interface. Colors indicate the VOC for which measurements were made. Bars indicate standard error of the mean. All labels use residues found in the wt system. Correlations were calculated using the Dynamical Network Analysis python package, and only contacts that had a mean correlation of 0.2 in at least one system were kept for this analysis.

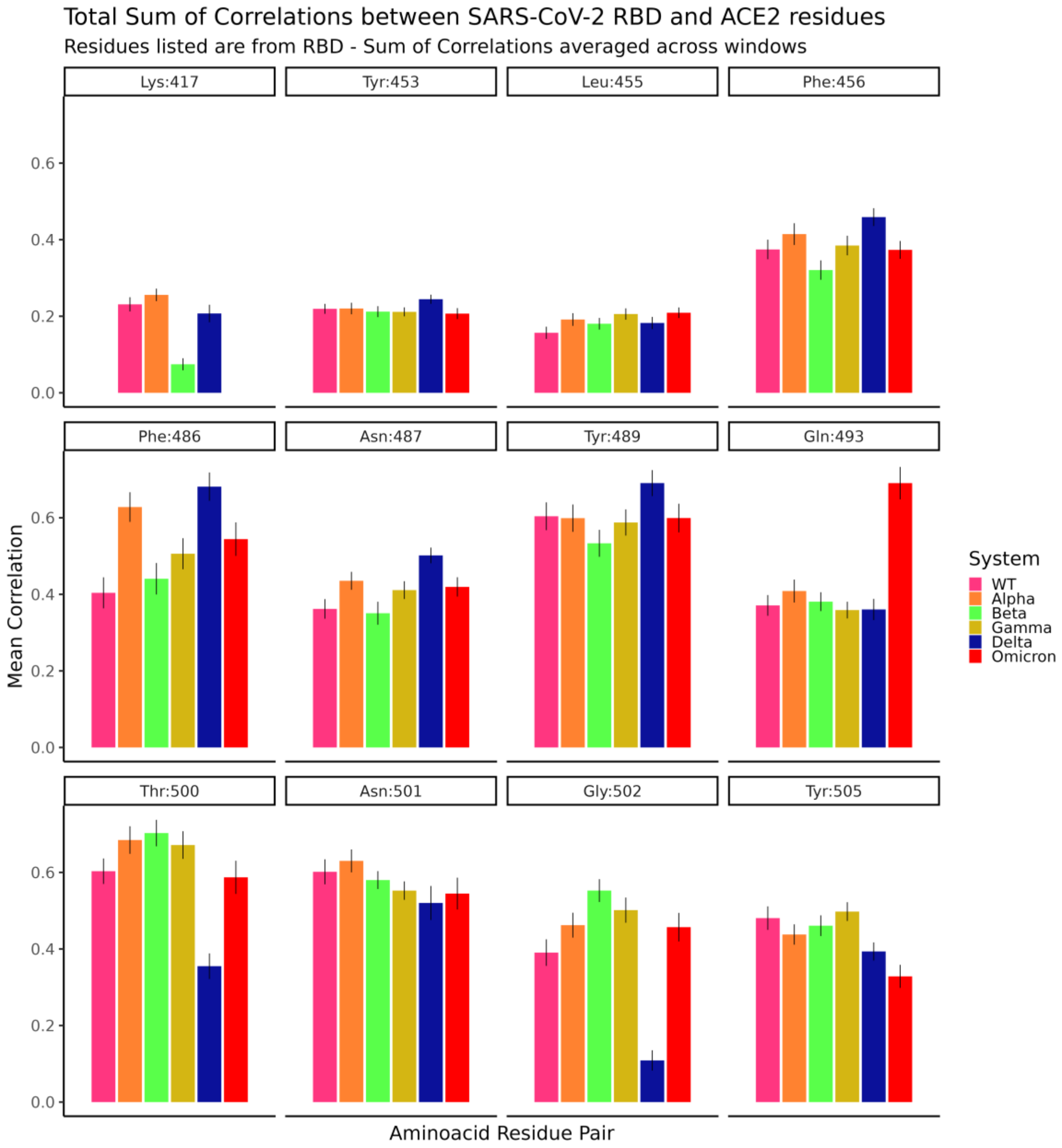


**Supporting Fig. S9.** Generalized correlation between the motions of residue pairs across the RBD:ACE2 interface. All correlations for each RBD residue were summed to indicate its overall contribution for interface stability. Bars indicate standard error of the mean. Colors indicate the VOC for which measurements were made. All labels use residues found in the wt system.

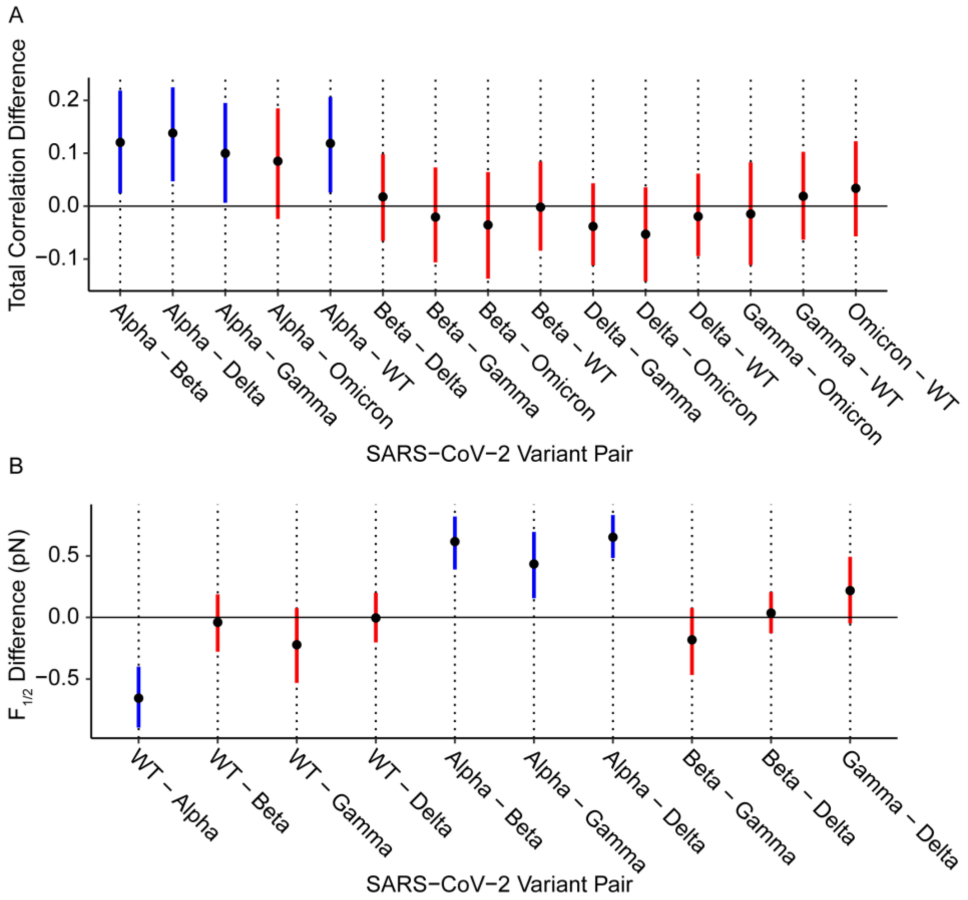


**Supplementary Fig. S10. Bootstrap analysis of Total Correlation and $F_{1/2}$. A** Confidence Intervals for the difference between the Total Correlation measurements across RBD:ACE2 interfaces. A CI that includes zero indicates the difference is not significant (in red). For VOC pairs with significant differences (in blue), the *p*-value was estimated using bootstrapping. For Alpha:Beta, $p = 0.04$; Alpha:Delta, $p = 0.02$; Alpha:Gamma, $p = 0.08$; For Alpha:WT, $p = 0.04$. **B** Confidence Interval for the difference between the $F_{1/2}$ measurements made for VOCs. P-value estimates were: For WT:Alpha, $p = 10^{-3}$; Alpha:Beta, $p = 10^{-3}$; Alpha:Gamma, $p = 0.03$; Alpha:Delta, $p = 10^{-3}$.
